# Crowding-induced phase separation and solidification by co-condensation of PEG in NPM1-rRNA condensates

**DOI:** 10.1101/2022.07.29.502035

**Authors:** Alain A.M. André, N. Amy Yewdall, Evan Spruijt

## Abstract

The crowdedness of the cell calls for adequate intracellular organization. Biomolecular condensates, formed by liquid-liquid phase separation of intrinsically disordered proteins and nucleic acids, are important organizers of cellular fluids. To underpin the molecular mechanisms of protein condensation, cell-free studies are often used where the role of crowding is not investigated in detail. Here, we investigate the effects of macromolecular crowding on the formation and material properties of a model heterotypic biomolecular condensate, consisting of nucleophosmin (NPM1) and ribosomal RNA (rRNA). We studied the effect of the macromolecular crowding agent PEG, which is often considered an inert crowding agent. We observed that PEG could induce both homotypic and heterotypic phase separation of NPM1 and NPM1-rRNA, respectively. Crowding increases the condensed concentration of NPM1 and decreases its equilibrium dilute phase concentration, while no significant change in the concentration of rRNA in the dilute phase was observed. Interestingly, the crowder itself is concentrated in the condensates, suggesting that co-condensation rather than excluded volume interactions underlie the enhanced phase separation by PEG. Fluorescence recovery after photobleaching (FRAP) measurements indicated that both NPM1 and rRNA become immobile at high PEG concentrations, indicative of a liquid-to-gel transition. Together, these results shed new light onto the role of synthetic crowding agents in phase separation, and demonstrate that condensate properties determined *in vitro* depend strongly on the addition of crowding agents.

**STATEMENT OF SIGNIFICANCE:** Liquid-liquid phase separation of proteins and nucleic acids leads to the formation of biomolecular condensates. To mimic biomolecular condensates in vitro, polymeric crowding agents, such as PEG, are often added. Such crowding agents are considered to make in vitro solutions more physiologically relevant, by mimicking the high cellular macromolecule concentrations. However, these crowding agents are commonly selected for their commercial availability and solubility in water, and their influence on phase separation and the physicochemical properties of condensates are seldom studied. Here we use biophysical methods to show that PEG induces phase separation of a model condensate through co-condensation rather than volume exclusion. As a consequence, crowding changes the partitioning, concentrations and viscoelastic properties of the condensates significantly, which sheds new light onto studies aimed at quantifying the material properties of biomolecular condensates.

## INTRODUCTION

One of the most fascinating aspects of the inner biochemistry of a cell is how it functions in an extremely crowded environment. Typical estimates indicate that cellular biopolymers, including proteins and RNA, occupy 20 to 30 vol% of the cell, limiting intracellular diffusion and making the cell’s interior a very crowded place (1–3). To enable effective biomolecular reactions, the cellular organization is of great importance (4,5). Traditionally, membrane-bound organelles have been extensively studied for their roles in organizing biochemical processes. This research has recently entered a new phase, as biomolecular condensates, also referred to as membraneless organelles, were found to be involved in the regulation of several biological processes, including transcription (6–8), cell signalling (9,10) and ribosome biogenesis (11–13).

Biomolecular condensates benefit from the absence of a physical membrane barrier, giving these condensates dynamic properties: they can fuse, ripen and wet membranes or become engulfed by other condensates (14). This reflects the molecular nature of these condensates: they are typically enriched in proteins containing intrinsically disordered regions (IDRs) and nucleic acids (11,15). The IDRs are not only disordered, they also contain certain repetitive motifs, for example in the form of weakly charged patches (16,17). Condensation is typically driven by liquid-liquid phase separation (LLPS) of these sticky moieties in the IDRs, but it is not limited to IDRs, as structural domains have been found to act as effective stickers as well (18–20). The balance between intermolecular association being not too strong to avoid turning the condensate into a gel, and not too weak to avoid dissolving it, makes most condensates highly responsive to changes in their environment, such as salinity and crowding, or subtle changes to the molecular constituents, such as enzymatic product formation or post-translational modifications (4,21).

Many of the molecular mechanisms underlying condensate formation and phase behavior have been unravelled through *in vitro* (here: cell-free) experiments (22–25). However, the cell-free environment in which proteins are studied is often far from realistic intracellular conditions. Factors that are typically regulated in what are considered to be ‘physiological’ conditions include ionic strength, pH and temperature (26). In contrast, the high degree of crowding in a living cell is not commonly taken into account in cell-free experiments despite the fact that many studies have shown that crowding can have a significant effect on protein stability, complexation and reactivity (3,4,27,28). When crowding is taken into account, the crowded cellular milieu is often mimicked by the addition of water-soluble polymers such as poly(ethylene glycol) (PEG), Ficoll and dextran (29–34). Although these polymers are highly water soluble, PEG and dextran can also undergo segregative phase separation, ending up in separate phases (35,36). This indicates that they have non-negligible interactions with each other, and suggests that these and other crowding agents could also interact with disordered and structured biomolecules, and affect their phase behavior. Therefore, a systematic study of the effect of crowding agents on commonly studied biomolecular condensates is of great relevance to understand the role of crowding in LLPS.

Crowding agents can affect biomolecular condensates in three ways. (i) They could promote (or in theory also suppress) phase separation by enhancing the weak intermolecular interactions through excluded volume effects or co-condensation, thereby shifting the binodal line to lower concentrations (37). (ii) Crowding agents can co-localize into the dense phase, as has been observed for dextran (38). This may be caused by the distinct local chemical environment (for instance nuage bodies are thought to have more hydrophobic interior (22)), but the change in composition could also lead to a further change in the local environment. (iii) Finally, the enhanced intermolecular interactions and altered composition could result in a change of the biophysical properties. For example, the condensed phase could become more viscous or switch to a solid-like state, such as previously observed for FUS (39) and NPM1 (29).

Here, we investigate the presence of these three effects in a well-studied condensate model of NPM1-rRNA using the most prevalent crowding agent found in studies of protein phase separation, PEG. We show that PEG induces both homotypic and heterotypic phase separation, and quantified the changes in the dilute and condensed phase. We observed the strongest crowding effect for NPM1, while there was no significant effect on rRNA in the dilute phase. Through confocal microscopy we were able to prove that PEG weakly partitions into the condensed phase, where it causes a differential increase in the NPM1 and rRNA density, thereby altering the condensate composition. Finally, using fluorescence recovery after photobleaching (FRAP) we measured the viscoelastic properties of both NPM1 and rRNA, which showed a rapid decrease of the mobile fraction in the presence of PEG.

## MATERIALS AND METHODS

### Materials

Unless otherwise stated all chemical were purchased from Sigma-Aldrich. All aqueous solutions were prepared in Milli-Q water (18.2 MΩcm), except for the rRNA stock solution which was dissolved in nuclease free water (Invitrogen).

### Protein expression and purification

*E.coli* BL21 (DE3) were transformed with pET28a(+)NPM1-wt. Bacterial cell cultures were grown in Luria Broth (LB-broth) supplemented with 50 μg L^-1^ kanamycin at 37 °C till OD_600_ reached 0.6-0.8 before expression was induced with 1 mM IPTG. Protein expression was carried out overnight at 18 °C and pelleted through centrifugation. Pellets were either stored at −80°C or directly used for purification. For purification, pellets were thawed on ice and resuspended in lysis buffer (10 mM Tris-HCl, pH 7.5, 300 mM NaCl, 20 mM imidazole) supplemented with 5 mM β-mercaptoethanol, 1× protease inhibitor (Roche), and 10 mM PMSF. Cell suspensions were either lysed by sonication (Sanyo Soniprep 500) or French Press homogenizer (Homogenizing Systems LTD). The lysate was cleared through centrifugation at 20,000 *g* at 4 °C for 30 minutes in a Beckman JA25.50 rotor. The supernatant was loaded on an equilibrated 5 mL His-trap column (GE healthcare/Cytiva) at 4 °C. The loaded column was washed with 10 column volumes (CV) lysis buffer and eluted with 3 CV elution buffer (20 mM Tris-HCl, pH 7.5, 300 mM NaCl, 5 mM β-mercaptoethanol, 500 mM imidazole). Protein containing fractions were dialysed overnight against SEC-buffer (10 mM Tris-HCl, pH 7.5, 300 mM NaCl, 1 mM DTT) and concentrated to <5 mL. The concentrated protein sample was loaded on a Superdex 200 16/600 (GE-healthcare) size exclusion column connected to an AKTA Basic FPLC (GE Healthcare) in SEC-buffer. Elution was carried out at room temperature at 1 mL/min and monitored at 205 nm, 254 nm and 280 nm. Fractions of the main peak were pooled, and concentrated. The concentration was determined using the NanoDrop One^C^ (Thermo Scientific), and aliquots were snap frozen in liquid nitrogen and stored at −80 °C.

### NPM1-Alexa488 labelling

NPM1-wt was labelled using AlexaFluor488 C5 maleimide dye (Thermo Fisher) according manufacturer’s protocol. In short, 100 μM NPM1 was dialysed against 10 mM Tris-HCl, pH 7.5, 300 mM NaCl, 1 mM TCEP. Using a Millipore spin filter (MWCO: 10 kDa), the excess TCEP was removed, and 200 μM AlexaFluor488 C5 maleimide dye was added, and incubated overnight at 4 °C. Excess dye was removed through dialysis (Millipore, MWCO 3.5 kDa) against SEC buffer, and the concentration was determined using the NanoDrop One^C^.

### Ribosomal RNA isolation

*E. coli* (DE3) pLysS cells were grown at 37 °C in LB-broth till OD_600_ reached 1.2, and pelleted through centrifugation (5,000 *g* at 4 °C for 15 minutes). Pellets were washed twice in S30 buffer A (50 mM Tris-HCl, pH 7.7, 60 mM potassium glutamate, 14 mM magnesium glutamate, 2 mM DTT), and resuspended in S30 buffer A (1 mL buffer to 1 gram cell pellet). This cell suspension was then homogenized through sonication (Sanyo Soniprep 150) and cleared through centrifugation (15,000 rpm at 4 °C for 25 minutes) in a Beckman JA25.50 rotor. Ribosomes were isolated by ultracentrifugation for 3 hours at 50,000 rpm (Beckman-Coulter Optima-90, with a fixed angle 90-Ti rotor). The glassy rough ribosome pellets were dissolved overnight in S30 buffer B (5 mM Tris-HCl, pH 8.2, 60 mM potassium glutamate, 14 mM magnesium glutamate, 2 mM DTT) at 4 °C. The ribosomal RNA (rRNA) was isolated from the ribosomes through standard phenol chloroform extraction using phenol:chloroform:isoamyl alcohol (PCI, 49.5:49.5:1). The final rRNA concentration was determined using the NanoDrop One^C^, where 1 OD_600_ = 40 μg mL^-1^ RNA and stored at −80 °C.

### RNA-Alexa647 labelling

The ‘3-end of the rRNA was labelled with AlexaFluor647-hydrazide using a periodate oxidation reaction (40–42). In short, to 80 μL of rRNA (3.4 mg mL^-1^), 7 μL of nuclease free water, 3.33 μL of 3 M sodium acetate (pH 5.2), and 10 μL of 25 mM sodium periodate (freshly prepared in water on the day) was added. The mixture was incubated on ice for 50 minutes. Subsequently, 20 μL of 3 M sodium acetate (pH 5.2) and 80 μL nuclease free water was added. The activated RNA was then precipitated by addition of 400 μL isopropanol through cooling it on ice for at least one hour. The RNA was then spun down (14,000 *g* at 4 °C for 15 minutes). The supernatant was removed and 150 μL of ice-cold ethanol was added to the pellet without mixing. After another centrifugation step, and removal of the supernatant, the RNA was mixed into the reaction buffer (100 mM sodium acetate, pH 5.2, 25 nmol Alexa-647 hydrazide). The reaction was left over 48 hours, after which the labelled RNA was isolated through a isopropanol and ethanol precipitation. The rRNA-A647 was then redissolved in 80 μL of nuclease free water. The concentration was determined using the NanoDrop One^C^.

### Preparation of NPM1-rRNA condensates

NPM1-AlexaFluor488 (NPM1-A488) stock solutions were prepared at 200 μM with 1:9 molar ratio AlexaFluor488 labelled NPM1 to unlabelled NPM1 in 10 mM Tris-HCl (pH 7.5), 300 mM NaCl. NPM1-A488 aliquots of 20 μL were snap frozen and stored at −80 °C.

The order of components for a typical experiment consists of first mixing the PEG (10 kDa, from a 35 wt% stock), with the buffer (from a 4x stock of 40 mM Tris-HCl (pH 7.5), 600 mM NaCl) and then diluted to the required final volume, often 30 μL at room temperature. 15 minutes prior to measuring the NPM1-A488 and rRNA were added to the premixed diluted buffer.

### Quantification of the dilute phase

A typical sample of 30 μL was prepared in 10 mM Tris (pH 7.5), 150 mM NaCl with varying concentrations of PEG, NPM1/NPM1-A488 (1:9 molar ratio labelled), and rRNA-A647 (only labelled) as described above. After incubating for 15-20 minutes at room temperature, the condensed phase was separated from the dilute phase by centrifugation at 21,000 *g* for 20 minutes at room temperature. The dilute phase was then transferred to a 384-well plate (Nunc, flat bottom) and the fluorescence intensity was measured on a plate reader (Tecan Spark M10) at 485/535 nm for NPM1-A488, and 620/680 nm for rRNA-A647. Concentrations of the dilute phase were calculated based on calibration curves (Fig. S1).

### Preparation of modified glass coverslips

Ibidi 18-well chambered slides (#1.5) were first cleaned using oxygen plasma and directly afterwards incubated in a 0.01 mg/mL solution of PLL(2)-g[3,5]-PEG(2) (SuSoS AG, Switzerland) in 10 mM HEPES buffer (pH 7.4) for at least 1 hour at room temperature. Ibidi chambers were rinsed several times with MilliQ water and dried with pressurised air. Modified glass slides were stored at −20 °C.

### Quantification of the condensed phase by confocal microscopy

Images for partitioning were acquired on a Leica Sp8x confocal inverted microscope (Leica Microsystems, Germany) equipped with a DMi8 CS motorized stage, a pulsed white light laser, and 2 x HyD SP GaAsP and 2x PMT detectors. Images were recorded using the LAS X v.3.5 acquisition software, using a HC PL APO 100x/1.40 oil immersion objective.

Samples were prepared as described above. After incubation of NPM1-rRNA condensates for 15 minutes, the sample was transferred to the modified Ibidi-18 well chambers and the condensates were allowed to settle to the bottom of the chamber for 10 minutes. Partitioning coefficients were analysed by MATLAB, the background was subtracted by measuring non-fluorescent condensates at the same settings as for the fluorescent images. The partitioning coefficient was then calculated using: *K_p_* = *(I_condensate_ – I_background_)/ (I_dilute_ – I_background_)* where *I_condensate_* is the average intensity of all condensates in one frame and *I_dilute_* the average intensity of the area without condensates. Standard deviations were determined of at least three sets of *K_p_* values derived from three different images.

### FRAP analysis

For FRAP analysis time lapse videos were recorded at room temperature on a CSU X-1 Yokogawa spinning disc confocal unit connected an Olympus IX81 inverted microscope, using a x100 piezo-driven oil immersion objective (NA 1.3) and a 488, 561 or 640 nm laser beams. Emission was measured with a 200 ms exposure time at a rate of 120 frames per minute, using an Andor iXon3 EM-CCD camera. The acquired images have a pixel size of 141 nm and a field of 72×72 μm^2^. For bleaching, a small region of interest (ROI) was selected in the middle of a condensed droplet. The 488 nm or 640 nm laser line was set to at 100% laser power using 75 pulses of 200 μs. The recovery was then imaged at reduced laser intensity with a time interval of 500 ms.

Recovery profiles were analysed using ImageJ, the normalized intensities were fitted to a 2D-diffusion with a fixed boundary (43). Using the resulted exponential decay equation *I_normalized_* = *A(1-e^-bt^)+C*, from which we obtained the parameters *A*, *b* and *C*. The recovery half-life was then determined at *t_1/2_*=*ln(2)/b*.

## RESULTS AND DISCUSSION

### PEG shifts the phase diagram of NPM1-rRNA

Inspired by the numerous membraneless organelles that contain both proteins and RNA, we chose a heterotypic system consisting of nucleophosmin (NPM1) and ribosomal RNA (rRNA) as model condensate. This system is illustrated in Fig. 1A, and forms liquid droplets under ‘physiological’ conditions (here defined as physiological salt by 10 mM Tris, 150 mM NaCl), even without crowding. By labelling both NPM1 and rRNA with fluorophores we could observe that indeed the rRNA co-localizes into the dense NPM1 droplets (Fig. 1B), in agreement with our previous study (40). We found a maximum degree of phase separation, as inferred from microscopy analysis, at a NPM1:rRNA ratio of 20 μM NPM1: 150 ng μL^-1^ rRNA (19,40). This corresponds roughly to one pentamer NPM1 interacting with 110 nucleotides of rRNA. When keeping this ratio constant, we could decrease the protein concentration to a lower limit of 10 μM and still observe liquid condensates (Fig.1C, S2).

**Figure 1:**
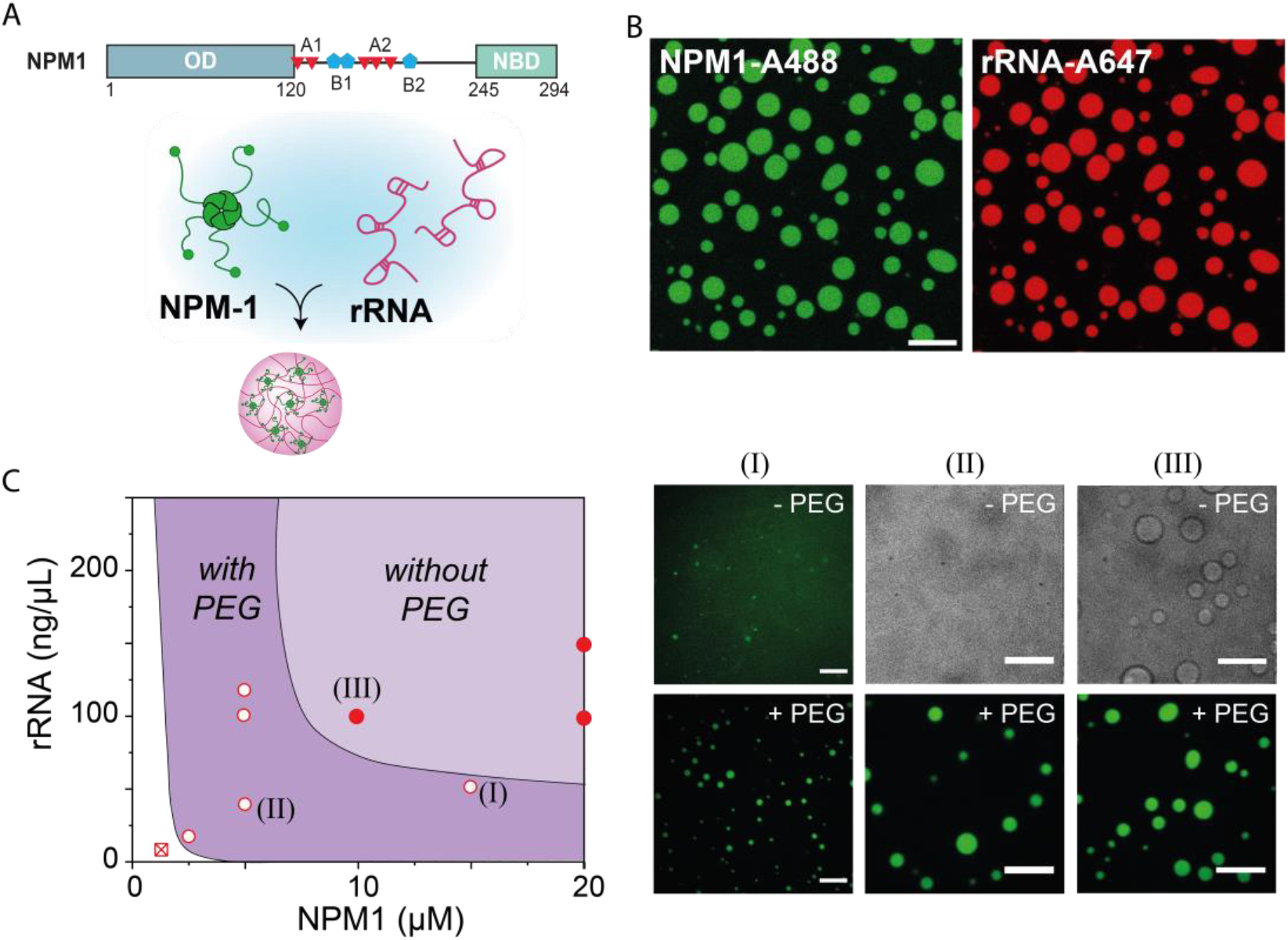
PEG induced phase separation of NPM1-rRNA. (A) Schematic illustration of nucleophosmin (NPM1) protein domains and the formation of condensates with rRNA. NPM1 encodes for a structural N-terminal oligomerization domain (OD) and a C-terminal nuclear binding domain (NBD) linked by an intrinsically disordered region with two acidic tracts (A1 and A2) and two weak basic tracts (B2 and B2). Pentamers of NPM1 phase separate with rRNA into condensates in vitro. (B) Fluorescent microscopy images of NPM1-rRNA condensates at 10 μM NPM1-Alexa488; 150ng μL^-1^ rRNA-Alexa647 (in 10 mM Tris pH 7.5, 150 mM NaCl). Scale bar, 10 μm. (C) Schematic representation of the shift in phase diagram of NPM1-rRNA liquid-liquid phase separation, and three conditions in the presence or absence of 2 wt% PEG: (I) 15 μM NPM1, 50ng μL^-1^ rRNA, (II) 5 μM NPM1, 37.5 ng μL^-1^ rRNA, (III) 10 μM NPM1, 75ng μL^-1^ rRNA. Scale bars, 10 μm.

When we added polyethylene glycol (PEG), a commonly used crowding agent, to the mixtures of NPM1 and rRNA, we observed phase separation at lower NPM1 concentrations. For 2 wt% PEG, we could observe phase separation for concentrations down to 2.5 μM NPM1 and 19 ng μL^-1^ rRNA (Fig. 1C, S2), which suggests that PEG enhances the association between protein and RNA. Moreover, we found that at 2 wt% PEG, NPM1 could also phase separate without RNA at 10 μM protein concentrations and higher into apparently homotypic droplets (Fig. 1C, S3). These findings are in agreement with a previous study of Kriwacki and co-workers, who showed that no second component, such as arginine-rich peptides or rRNA, is needed for phase separation of NPM1 under PEG-based crowding conditions (29,30). These findings suggest that crowding by PEG not only enhances associative interactions between NPM1 and rRNA, but it enhances the self-interactions of NPM1 even more. Assuming that volume exclusion by the crowding agents is responsible for enhancing the intermolecular interactions, the observed effects could be explained by the structural differences between NPM1 and rRNA: the protein NPM1 has a more globular shape and a larger effective radius than the polymeric rRNA and will therefore experience stronger volume exclusion due to crowders. Crowding by PEG will thus likely affect the concentrations of NPM1 and rRNA in both the dilute and the dense (condensate) phase in a non-trivial way. Therefore, we next sought to quantify the NPM1-rRNA phase diagram under crowding conditions.

### PEG reduces only NPM1 concentrations in the dilute phase

Excluded volume theory predicts that crowding enhances the effective attractions between macromolecules by increasing the entropy of the crowders upon complexation of the macromolecules. For our phase separating system, enhanced attraction could result in phase separation at lower concentrations, as suggested by the measurements in Figure 1C. To obtain a quantitative understanding of the effect of crowding on phase separation between NPM1 and rRNA, and how this affects their concentrations in the dilute and condensed phase, we determined the compositions of the droplets and supernatant as function of the amount of crowding agent PEG.

First, we examined the concentrations of NPM1 and rRNA in the dilute phase. The condensates were separated from the dilute phase by centrifugation (Fig. S4). We quantified the concentration of NPM1 and rRNA in the dilute phase for three different overall mixing ratios. In all cases, the addition of PEG reduced the concentration of NPM1 in the dilute phase, while the rRNA remained approximately constant or even increased (Fig. 2A). It is clear that this is not the expected behavior for a classical crowding agent that enhances the attraction between NPM1 and rRNA. In that case, the saturation concentration of both NPM1 and rRNA should decrease with increased crowding, although the relative degree of decrease might depend on the overall mixing ratio. Instead, we found that the rRNA saturation concentration remained constant for all mixing ratios. Crowding thus seems to leave the interaction between NPM1 and rRNA unchanged, while it enhances the interactions of NPM1 with itself, as the saturation concentration of NPM1 decreased significantly. Our results regarding the NPM1 concentrations in the dilute phase corroborate what has previously been observed for homotypic NPM1 condensation (29): crowding decreased the solubility of NPM1. However, it was also observed that the ratio between the arginine-rich SURF6 peptide (S6N) and NPM1 remained constant with increasing crowding concentrations, probably because the short peptide partitions as a client into the homotypic NPM1 droplets.

**Figure 2:**
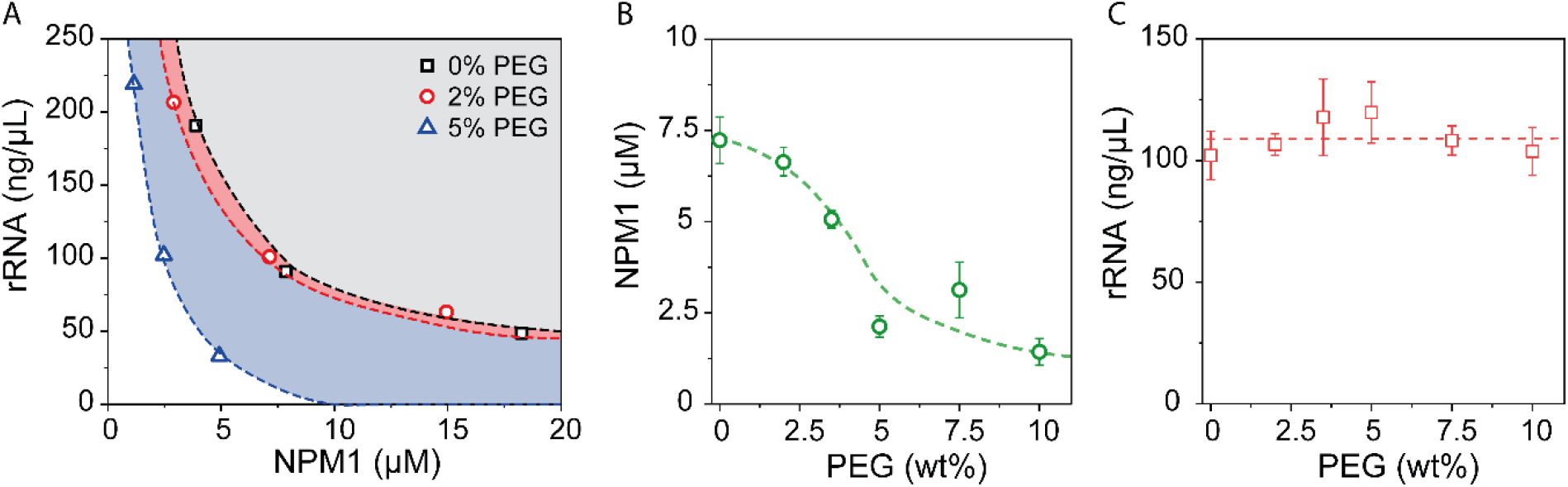
PEG reduces NPM1 concentration and not the rRNA concentrations in the dilute phase. (A) Concentrations of NPM1 plotted against rRNA in the dilute phase for three fixed NPM1:rRNA ratios: 5:150, 10:100, 20:50 (μM NPM1: ng μL^-1^ rRNA), at three different PEG concentrations. (B-C) Addition of PEG to 10 μM NPM1, 100 ng μL^-1^ rRNA reduces the concentration of NPM1 in the dilute phase (B) but not for rRNA (C). The errors in these figures are standard deviations from triplicate measurements, and the dashed lines are present to guide the eye.

To analyze the effect of PEG in more detail, we focussed on one ratio of NPM1:rRNA (10 μM NPM1 with 100 ng μL^-1^ rRNA), and looked at several concentrations of PEG (Fig. 2B and C). The dilute phase NPM1 concentration gradually decreased from 7.5 μM without crowding to 2.5 μM NPM1 at 10 wt% PEG (Fig. 2B). We did not observe a plateau at high crowding, despite the fact that the condensates had turned into gel-like structures with very little relaxation already at 2 wt% PEG (see also section: Crowding reduces condensate fluidity), but rather a gradual decrease that becomes asymptotic towards zero, in agreement with simple theoretical predictions for crowding-induced phase separation. For rRNA, we observed no clear change in the concentration in the dilute phase, as was found for other mixing ratios as well: it remained constant around 100 ng μL^-1^ (Fig. 2C).

### PEG partitions into NPM1-rRNA condensates and increase local concentrations

We next analyzed how crowding affects the composition of condensates by studying the condensed phase in more detail. The tiny combined volume of the condensate droplets (< 0.1 μL) made an equivalent analysis of the absolute concentrations by fluorescence spectroscopy impossible. We therefore analyzed the relative changes in the composition of the condensates using fluorescence microscopy (Fig. 3A), despite the intrinsic limitations of quantitative fluorescence at high concentrations (44,45). Here, we used labelled NPM1 and rRNA at low enough concentrations that the fluorophores are further apart than their typical homo-FRET distance (46).

**Figure 3:**
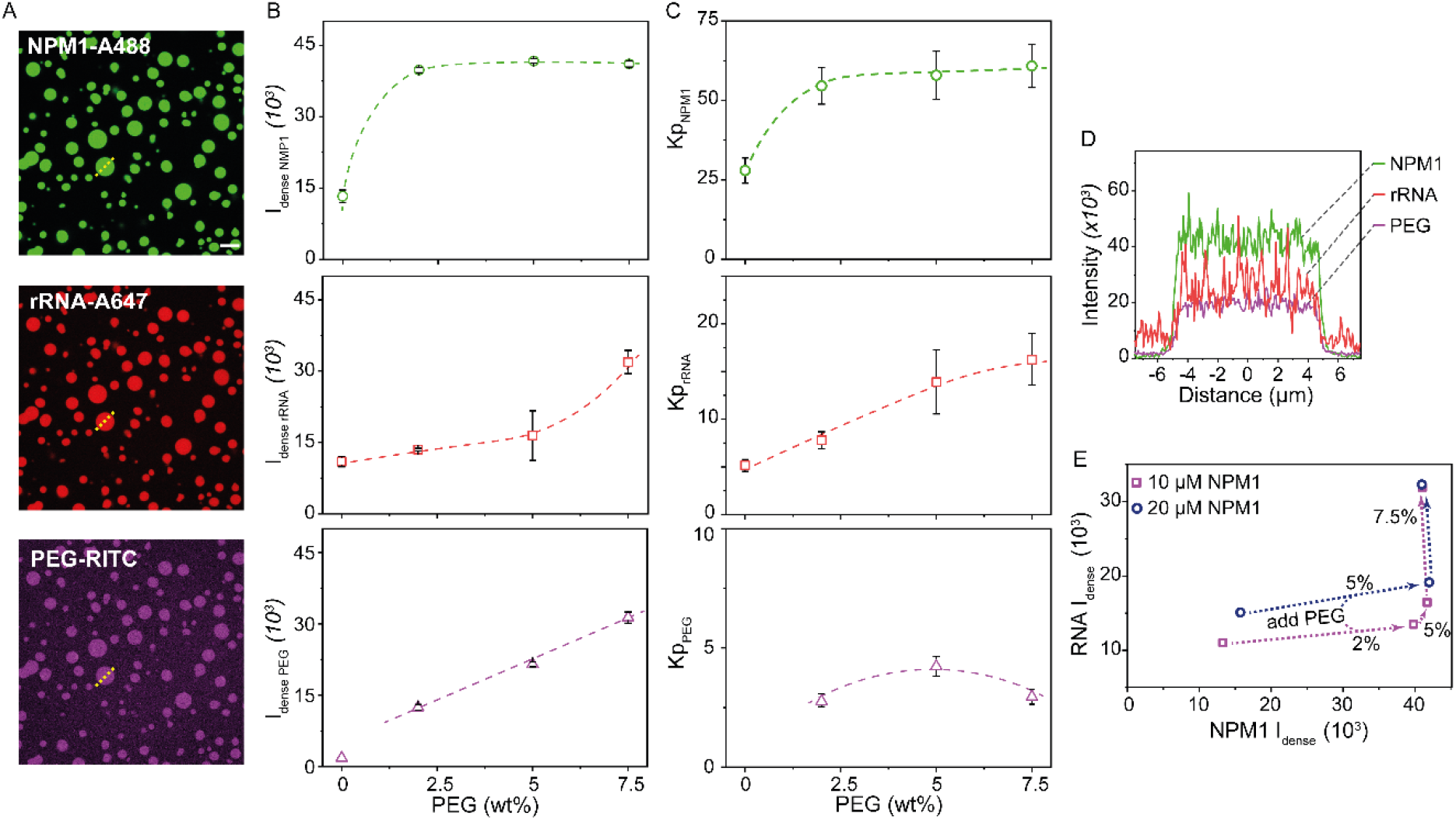
Confocal microscopy analysis of NPM1-rRNA condensates under crowded conditions. (A) Fluorescence images of NPM1-Alexa488, rRNA-Alexa647, and PEG-rhodamine, indicating PEG co-localizes into the condensed phase (20 μM NPM1, 150 ng μL^-1^ rRNA, 5 wt% PEG). Scale bar, 10 μm. (B-C) Average fluorescence intensities (B) and partitioning coefficients (K_p_) (C) of NPM1, rRNA, and PEG plotted against the concentrations of PEG. (D) Intensities profile across one single condensate (dashed line in A). (E) Graph representing the influence of crowding on the intensities (and therefore concentration) of NPM1 and rRNA within condensates. The errors in this figure are standard deviations from triplicate measurements, and all dotted lines and arrows are to guide the eye.

Confocal microscopy images show a three-fold increase in fluorescence intensity of NPM1 (Fig. 3B) upon addition of 2 wt% PEG. The partitioning coefficient (K_p_) was determined based on this fluorescence, and as a consequence the K_p_ for NPM1 (Fig. 3C) is showing the same trend. Further increasing the concentration of PEG did not lead to additional increase in NPM1 fluorescence, and most likely in concentration. An increase in NPM1 concentration upon crowding is expected if the crowders enhance the association between NPM1 proteins, although the magnitude of the observed increase is higher than expected for a crowder at approximately 5 vol% (47). Moreover, in the case of pure excluded volume interactions between crowder and the phase separating biomolecules, the local concentration of biomolecules inside the condensates and thereby their density is expected to continue increasing when the crowder concentration is increased further, since increasing the crowder concentration leads to further enhancement of the association strength between the biomolecules, which translates into a higher binodal condensate concentration. The fact that we do not observe a further increase in density of the NPM1 concentration inside the condensates suggests that the interactions between NPM1 and PEG may not be limited to volume exclusion.

For rRNA, we found that addition of PEG did not affect the concentration in the dilute phase. In general, rRNA showed a much lower partitioning into the condensates (K_p_ = 5) than NPM1 in the absence of PEG, and the rRNA concentration increased only slightly upon addition of PEG (Fig. 3B). At low (0-5 wt%) PEG concentration, the rRNA concentration increased about 25%, whereas the local NPM1 concentration increased threefold. However, at the highest PEG concentration (7.5 wt%), the rRNA concentration and partitioning increased about twofold, while the NPM1 concentration did not increase further at this PEG concentration (Fig. 3E), and also the rRNA concentration in the dilute phase did not change. The slight increase in rRNA in the condensed phase at low PEG concentrations is in agreement with our observation that the dilute phase concentration did not increase: PEG mostly enhances NPM1 self-interaction, which causes the condensed phase to become denser, but the interaction between NPM1 and rRNA is not strongly affected. Moreover, PEG is known to enhance folding of RNA into more compact states (48,49), which could explain the higher concentration of rRNA inside the condensates. The strong increase in rRNA concentration at the highest PEG concentration suggests that the condensates may no longer be pure liquids governed by LLPS at that point, but that they are kinetically trapped in a gel state, as we will discuss further below.

Finally, we also looked at the distribution of the crowding agent PEG over the two phases (Fig. 3). In the case of ideal excluded volume interactions linked to a first-order phase transition of a solute that is driven by a crowding-enhanced association, and ignoring the contribution of the crowders to the osmotic pressure, the concentration of crowders in the dilute and condensed phase is predicted to be the same. More generally, the high local concentration of biomolecules in the condensates exclude volume for the crowder molecules, and therefore, we expect that crowders are weakly depleted from the condensates. However, when we measured the distribution of PEG over both phases via fluorescence microscopy, we found that PEG was enriched in the condensed phase by a factor of 3 (Fig. 3B-C). To minimize the effect of the dye, less than 0.1% of the PEG was labelled in this study. Interestingly, the enrichment was independent of crowding concentration, and suggests that PEG exhibits associative interactions with NPM1 or rRNA or both. Therefore, the classical picture of PEG as an inert crowding agent that interacts via excluded volume interactions with biomolecules is not accurate. Instead, PEG seems to co-condense with NPM1 and/or rRNA in a form of ternary associative phase separation.

Partitioning of crowding agents was also observed by a few other groups. Hammer and co-workers observed similar partitioning of their small molecular weight dextran (4.4 kDa) in condensates made of the LAF1 RGG domain as in our PEG-based crowding studies (38). Moreover, they observed decreasing partitioning of dextran when its size was increased. In a systematic study on coacervates made of spermine and polyuridylic acid (polyU), Keating and co-workers observed a similar effect of PEG (8 kDa) and Ficoll (70 kDa) as crowding agents on the induction of phase separation. In contrast to our study, they observed a weak exclusion of PEG, even though its size was similar, but an enrichment of Ficoll into spermine-polyU coacervates (50). Their results indicated that for spermine-polyU coacervates, PEG behaves more like an inert crowder. However, in both cases (crowding agent exclusion and inclusion) a promotion of phase separation was observed, which they contributed to an enhancement of the favourable base-stacking of polyU. If the exclusion of PEG by polyU-based coacervates is taken as an indication that there is no specific association between PEG and RNA in the condensed phase, we can infer that in our system of NPM1-rRNA, PEG is mostly associated with the NPM1. This suggests that PEG-induced phase separation of homotypic NPM1 droplets is likely driven by a co-condensation of PEG and NPM1 (Fig. S3). The same may be the case for a significant number of other homo- and heterotypic IDP-based condensates that have been reported in the presence of PEG (4), for example a recent preprint by Knowles and co-workers indicated PEG interacts with FUS-protein condensates (51).

Finally, our results show that the ratio between NPM1 and rRNA in condensates is altered upon increasing PEG concentration, in contrast to the NPM1-S6N condensates by Kriwacki and co-workers, which rely on electrostatic interactions (29). Our results indicate that the co-condensation with PEG, driven by weak associations between the PEG and NPM1, is responsible for altering the condensate composition, and increasing the overall macromolecular concentration within the condensed phase. By extension, a similar mechanism may play a role in the crowded environment in the cell: besides excluded volume interactions, there may be many (weak) soft repulsive or attractive forces (3,52) between all the crowders and phase separating IDRs, which could result in promotion of phase separation through co-condensation or segregation (53). This might also lead to changes in the material properties of condensates formed in the presence of crowding agents, as we investigated next.

### Crowding reduces condensate fluidity

Liquid-liquid phase separation is characterized by the formation of liquid droplets that typically exhibit rapid recovery of fluorescence after photobleaching (FRAP). Indeed, our model system shows full and fast recovery after photobleaching in the absence of PEG (Fig. 4). The mobile fraction correlates to the recovery percentage, and for NPM1 in buffer this is almost 90%, and for rRNA around 65% (Fig. 4D). This indicates that NPM1 shows almost a full recovery, since the total recovery was not corrected for the size of bleached area. From the recovery curves we were able to determine the recovery half-time (t_1/2_), which was for both NPM1 and rRNA approximately 10 seconds. Thus, although NPM1 and rRNA have similar characteristic recovery times, a larger fraction of the rRNA in condensates was immobile compared the NPM1. We attribute this to possible RNA-RNA interactions (40).

**Figure 4:**
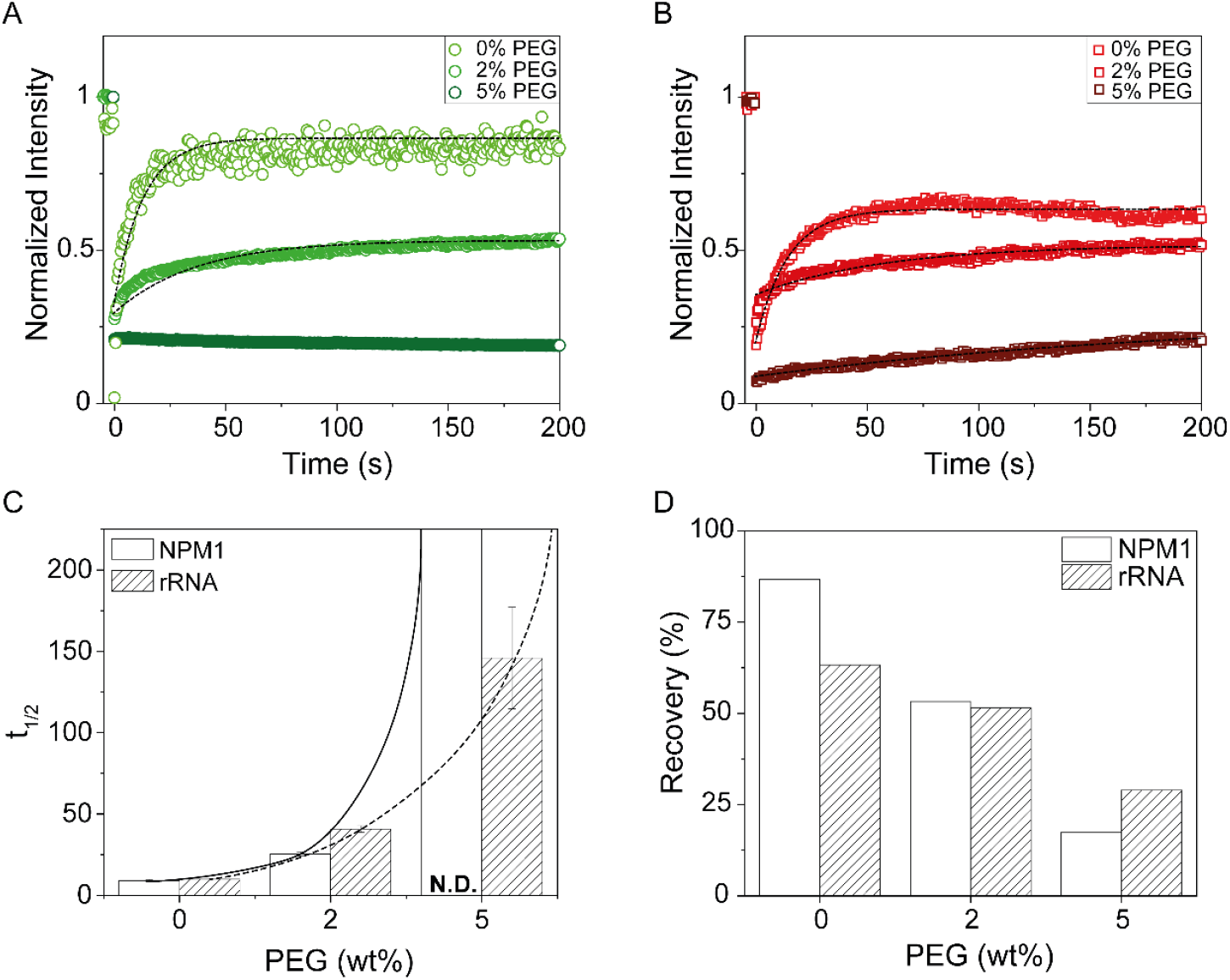
PEG changes the viscoelastic properties of NPM1 and rRNA. (A-B) FRAP recovery curves of NPM1 (A), and rRNA (B) upon increasing PEG crowding. The dashed lines represent the model fit for which the t_1/2_ and total recovery were determined. (C) Calculated half-life (t_1/2_) of NPM1 and rRNA at different concentrations of PEG. The lines are there to guide the eye. (D) Mobile fraction from the total recovery determined from the FRAP curves.

The addition of PEG reduced the mobile fraction of both NPM1 and rRNA in the condensed phase (Fig. 3) significantly: 2% PEG decreased the mobile fraction of both NPM1 and rRNA to 50%, while the recovery half-time increased to 25 seconds and 40 seconds respectively. When we increased the PEG concentration further to 5%, both NPM1 and rRNA became completely immobile, indicating that the condensates are no longer liquid droplets governed by LLPS, but have turned into gels.

The solidifying effect of PEG on both NPM1 and rRNA is interesting compared to our previous results. For instance, our lab showed that physiological concentrations of Mg^2+^ was only affecting the rRNA diffusion in NPM1-rRNA condensates (40) by enhancing interactions between the rRNA molecules. Kriwacki and co-workers demonstrated that the addition of the arginine rich domain SURF6 could liquefy homotypic NPM1-NPM1 condensates (29). These newly formed NPM1-SURF6 condensates depend on oppositely electrostatic interactions, while NPM1-rRNA condensation is considered to rely on the RNA recognition motifs (RRMs) present in NPM1.

Taking into account that PEG is present within the condensed phase (Fig. 3) we hypothesize that co-condensation of PEG is solidifying NPM1-rRNA droplets in vitro. Since PEG is not fulfilling the classical crowding model of an inert macromolecule, we observe that PEG likely binds to NPM1 without affecting the ability of NPM1 to bind rRNA via the RRMs. The co-condensation of PEG and NPM1 strongly increases the NPM1 content, but also reduces their diffusion. The rRNA remains condensed with the NPM1 via binding to the RRMs, but its concentration is not increased as much as NPM1 due to weakly unfavourable interactions with the PEG, as also suggested by the work of Keating and co-workers (50). Nevertheless, the higher local concentration of NPM1 also decreases the diffusivity of rRNA, as it is more likely to be bound by multiple RRMs of NPM1. Above a threshold PEG concentration, the local interactions between NPM1-NPM1 and NPM1-PEG become too strong and their mobility decreases sharply, which is reflected by the absence of NPM1 recovery in Fig. 4C.

## CONCLUSIONS

In conclusion, we reported that the crowding agent PEG could induce and enhance phase separation of a model biomolecular condensate consisting of NPM1-rRNA. By quantifying the compositions of both the condensed and dilute phase, we could deduce that only the protein component NPM1 is depleted from the dilute phase and enriched in the condensed phase. Although the concentration of rRNA remained constant in the dilute phase, the condensed phase showed a slight PEG-dependent enrichment, possibly due to a more condensed state of the dense phase. Through fluorescent labelling of the crowding agent, we found that, surprisingly, PEG is also enriched in the condensed phase, suggesting that it enhances phase separation by co-condensing with NPM1 rather than through excluded volume interactions. These results also indicate that partitioning of “crowding agents” can change condensate compositions and material properties, and that even crowders that are widely believed to be inert can have significant interactions with biomolecules that undergo liquid-liquid phase separation. This is relevant to the crowded environment of the cell as well: it is very likely that an important fraction of the macromolecules present in the cell exhibit weak repulsive or attractive interactions with the components of membraneless organelles, which results in altered composition and material properties compared to their in vitro reconstituted analogues. Finally, our results also put *in vitro* condensates formed in the presence of common crowding agents in a new perspective and suggest that addition of common crowding agents could affect *in vitro* phase separation systems and therefore should be selected with care.

## AUTHOR INFORMATION

### Author Contributions

Alain A.M. André: conceptualization, methodology, investigation, writing. N. Amy Yewdall: methodology. Evan Spruijt: conceptualization, software, writing, supervision funding acquisition.

## ACKNOWLEDGEMENTS

The authors thank Prof. Richard Kriwacki (St. Jude Children’s Research Hospital) for providing us with the NPM1-GFP plasmid, Aafke Jonker and Frank Nelissen (Radboud University) for providing protocols to isolate ribosomes and labelling RNA, Merlijn van Haren (Radboud University) for providing a MATLAB script to determine partitioning coefficients of the microscopy data and Dr. Ioannis Alexopoulos (Radboud University, now Justus Liebig University) for his guidance in confocal microscopy. We would like to thank Prof. Wilhelm Huck (Radboud University) and Prof. Allen Minton (NIH) for fruitful discussions. This work was financially supported by the Netherlands Organization for Scientific Research (NWO).

